# Genetic signatures of adaptation to sudden, extreme, and unprecedented environmental changes

**DOI:** 10.64898/2026.02.27.708566

**Authors:** Sebastian E. Ramos-Onsins, Luca Ferretti

## Abstract

Small natural populations are increasingly exposed to environmental shifts that are unprecedented in speed and intensity on population-genetics timescales. If populations persist, adaptation to these abrupt changes is expected to leave localized footprints of selection around the loci contributing to the adaptive response, in contrast to genome-wide signals produced by demography alone. Here we provide an explicit theoretical description of genetic footprints expected under rapid adaptation from standing genetic variation. We derive exact analytical results for how single-locus sweeps and polygenic shifts driven by sudden environmental changes distort the site-frequency spectrum (SFS) of linked mutations, and we link these distortions to the expected changes in standard diversity measures and neutrality statistics. Our results provide a guideline for detecting and interpreting genetic signatures of adaptation to extreme events, including those associated with climate change, particularly in small populations where adaptation or evolutionary rescue often proceed quickly and from preexisting variation.

## 1 Introduction

Contemporary climate change is not a single, smooth trend but a collection of sudden, extreme, and sometimes unprecedented environmental perturbations such as heat waves, droughts, marine heat events, altered seasonality, and novel combinations of stressors, all occurring at rates that can be rapid relative to the timescales over which populations historically adapted Grant *et al*. (2025). For many taxa, especially macroscopic eukaryotes with long generation times, low fecundity, and habitat fragmentation, such changes coincide with small census sizes and small effective population sizes Radchuk *et al*. (2019). These conditions reduce the efficacy of selection, increase the influence of drift, and elevate extinction risk Bellard *et al*. (2012).

When environmental change drives rapid population decline, genomic variation can be altered genome-wide through bottlenecks and increased drift Caplins *et al*. (2014). However, extremely fast demographic collapse can leave relatively weak or nonspecific genomic “footprints” beyond those expected under severe drift, because it acts broadly on all alleles and does not preferentially elevate particular haplotypes Peery *et al*. (2012). By contrast, when populations adapt to a novel environment, particularly in the context of evolutionary rescue when survival depends on a small set of traits, selection can increase the frequency of specific alleles (or coordinated sets of alleles), leaving localized signatures around the causal variants Stephan (2010).

A central challenge is to distinguish localized signals of selection associated with adaptive rescue from background demographic effects in small, vulnerable populations. This is complicated by three features common in the scenarios motivating this work:

- small effective population size implies weak efficacy of selection for modest-effect variants, and therefore adaptation driven by large-effect variants or by multiple variants at the same time;
- extreme environmental shifts can impose strong selection pressures over short intervals;
- limited time makes adaptation from *de novo* mutation unlikely in many cases, making standing genetic variation (and potentially introgression) the primary substrate for rapid response.

Here we develop an analytically tractable framework to characterize the genetic footprints of rapid adaptation from standing variation following sudden environmental change. We focus on the site-frequency spectrum of linked variation near selected loci and derive expected distortions under (i) strong selection at a single locus and (ii) polygenic adaptation where many loci undergo small frequency shifts. We then translate these distortions into predictions for common summary statistics (e.g. Watterson’s *θ*_*W*_ Watterson (1975), *nucleotide diversity π* Tajima (1983), *Tajima’s D* Tajima (1989), *Fay and Wu’s H* Fay and Wu (2000)).

## 2 Model and Methods

### 2.1 Site Frequency Spectrum statistics

*The frequency spectrum is the number of mutations of a given frequency in a sequence of length L*. We denote the frequency spectrum for a sample of size *n* by *ξ*_*k*_, where *k* = 1 … *n* − 1 is the derived allele count in the sample. The population spectrum is denoted by *ξ*(*f*), where *f* is the allele frequency.

The frequency spectrum statistics used in this paper are the Watterson estimator of variability (Watterson (1975)):

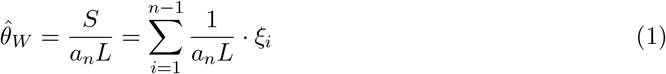

the pairwise nucleotide diversity (Tajima (1983)):

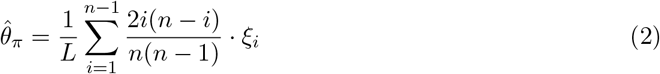

Tajima’s *D*, which measures the abundance of common versus rare alleles (Tajima (1989)):

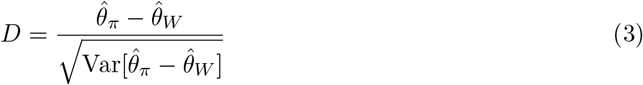

and Fay and Wu’s *H*, which tests for classical selective sweeps (Fay and Wu (2000)):

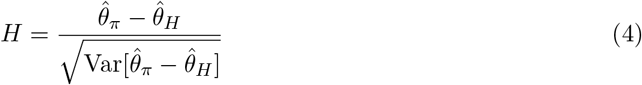

where 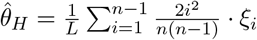

The standard neutral model results in the expected spectra 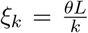 at the sample level and 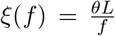 at the population level, where *θ* is the rescaled mutation rate per base. Under this model, the estimators above are unbiased (i.e. 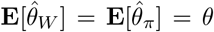) and the neutrality tests are centered (i.e. **E**[*D*] = **E**[*H*] = 0).

### 2.2 Model overview and assumptions

We consider a natural population that experiences an abrupt environmental shift at time *t* = 0, after which selection favors alleles that increase fitness under the new conditions. We study genetic variation in a genomic region closely linked to a focal selected variant (or, in the polygenic case, near one of many contributing variants). Our main object is the site-frequency spectrum (SFS), i.e. the distribution of derived allele frequencies among mutations in the linked region.

We emphasize regimes motivated by adaptation or evolutionary rescue in small populations:

- Adaptation proceeds primarily from standing variation present at *t* = 0.
- Selection is rapid and strong relative to drift over the short adaptive episode (with caveats discussed below).
- Recombination is ignored to obtain explicit closed-form expressions for the immediate linked signatures near the selected site; recombination is treated as a limitation and discussed in Section 4.

### 2.3 Wright-Fisher description of rapid selection from standing variation

We model allele-frequency changes at a selected locus in a discrete-generation Wright–Fisher framework. Let the favored allele have frequency *f*_0_ at the onset of the environmental shift and current frequency *f*_*now*_ after several generations of selection. Under strong, rapid selection, the stochastic contribution from drift during the adaptive episode may be small compared to the deterministic change imposed by selection, particularly when conditioning on the realized final frequency *f*_*now*_. In this regime, the short-term effect of selection on linked neutral variation can be described by a reweighting (or rescaling) of the two genetic backgrounds: chromosomes carrying the favored allele versus those carrying the alternative allele.

Intuitively, selection increases the representation of the favored background among sampled chromosomes. If recombination is negligible over the short interval and we condition on the final frequency of the selected allele, the process is equivalent to a single episode that “amplifies” the favored subpopulation and correspondingly “contracts” the alternative subpopulation, followed by sampling.

### 2.4 Derivation of the spectrum for partial sweeps under strong selection

We consider a population of constant effective size *N*_*e*_, containing a mutation that evolved neutrally until *t*_*s*_ generations ago. After that, the mutation was subject to strong selection pressure until present. This partial sweep brings the mutation from an initial frequency *f*_0_ in the population at time *t* = −*t*_*s*_ to a final frequency *f*_*now*_ at present (*t* = 0). We assume that the initial frequency *f*_0_ = *f* (−*t*_*s*_) and final frequencies *f*_*now*_ = *f* (0) of the derived allele are both not too small or too large, that is, 1*/N*_*e*_ ≪ *f*_0_, *f*_*now*_, 1 − *f*_0_, 1 − *f*_*now*_. Under strong selection, the sweep time *t*_*s*_ is very short with respect to coalescent times, i.e. *t*_*s*_ ≪ 2*N*_*e*_, therefore genetic drift has no time to cause a relevant change in the frequencies between *t* = *t*_*s*_ and *t* = 0 and selection behaves deterministically.

The mean frequency spectrum of the sequence before the sweep is described by the neutral frequency spectrum of linked sites *ξ*(*f* |*f*_0_) presented in Ferretti *et al*. (2018). There are several components in this spectrum, depending on their linkage to the focal mutation:

- strictly nested *ξ*^(*sn*)^: mutations occurring in a proper subset of the individuals with the focal mutation;
- co-occurring *ξ*^(*co*)^: mutations occurring on the same individuals as the focal mutation;
- enclosing *ξ*^(*en*)^: mutations that occur in all individuals with the focal mutation and more;
- complementary *ξ*^(*cm*)^: mutations occurring in all individuals without the focal one;
- strictly disjoint *ξ*^(*sd*)^: mutations present in a set of individuals that is non-overlapping and not complementary with the focal one.

The final spectrum depends on which allele (ancestral or derived) is positively selected. Both cases can be studied in this formalism: positive selection on the derived allele will correspond to *f*_*now*_ *> f*_0_ and positive selection on the ancestral allele will correspond to *f*_*now*_ *< f*_0_.

The effect of strong selection on the population spectrum is better understood by dividing the population in the two subpopulation corresponding to the two alleles of the focal mutation. The frequency of the alleles within each subpopulation remains constant during the sweep, while the relative size of the two subpopulations change from *f*_0_: 1 − *f*_0_ to *f*_*now*_: 1 − *f*_*now*_.

For example, a mutation of frequency *f* in the population that is “nested’ inside the focal mutation would have frequencies of *f/f*_0_ and 0*/*(1 − *f*_0_) inside the subpopulations; after the sweep, its final frequency in the population is *f*_*now*_(*f/f*_0_) + (1 − *f*_*now*_)(0*/*(1 − *f*_0_)) = *f*_*now*_*f/f*_0_. A similar approach can be applied to all the other components. We report the results for a linked mutation of initial frequency *f* in the following table.

| Component of the spectrum | Frequency in subpop linked to derived allele | Frequency in subpop linked to ancestral allele | Frequency $f_{final}$ in the final population |
| --- | --- | --- | --- |
| strictly nested | $f/f_0$ | 0 | $f_{now}f/f_0$ |
| co-occurring | 1 | 0 | $f_{now}$ |
| enclosing | 1 | $(f - f_0)/(1 - f_0)$ | $f_{now} + (1 - f_{now})(f - f_0)/(1 - f_0)$ |
| complementary | 0 | 1 | $1 - f_{now}$ |
| strictly disjoint | 0 | $f/(1 - f_0)$ | $(1 - f_{now})f/(1 - f_0)$ |

The new values for the components of the population spectrum can be obtained by transforming the densities through rescaling:

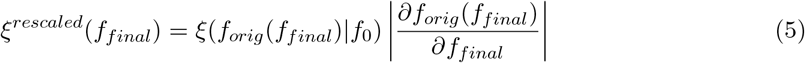

where the original frequency is expressed as a function of the final frequency *f* of the mutation as follows

| Component of the spectrum | Formula for original frequency $f_{orig}(f)$ |
| --- | --- |
| strictly nested | $f \cdot f_0/f_{now}$ |
| co-occurring | $f_0$ |
| enclosing | $f_0 + (f - f_{now})(1 - f_0)/(1 - f_{now})$ |
| complementary | $1 - f_0$ |
| strictly disjoint | $(1 - f_0)f/(1 - f_{now})$ |

Note that since some of these subspectra are densities, their rescaling involves a Jacobian transformation factor $J = |\partial f_{orig}(f_{final})/\partial f_{final}|$ , which can be extracted from the table above. These factors are $J_{(sn)} = f_0/f_{now}$ , $J_{(en)} = J_{(sd)} = (1 - f_0)/(1 - f_{now})$ .

Note that since some of these subspectra are densities, their rescaling involves a Jacobian transformation factor *J* = |∂*f*_*orig*_(*f*_*final*_)*/*∂*f*_*final*_|, which can be extracted from the table above. These factors are *J*_(*sn*)_ = *f*_0_*/f*_*now*_, *J*_(*en*)_ = *J*_(*sd*)_ = (1 − *f*_0_)*/*(1 − *f*_*now*_).

The final frequency *f*_*now*_ in the population may be unknown, but the final frequency in the sample *k*_*now*_*/n* ∼ *f*_*now*_ could be observed. Since a sample is a random choice of alleles from the population, the average spectrum is a convolution of the population spectrum and a binomial. The five components of the average spectrum for a sample are:

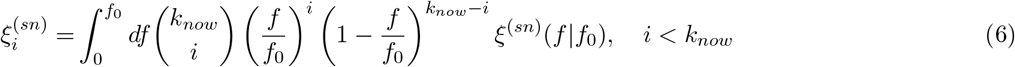

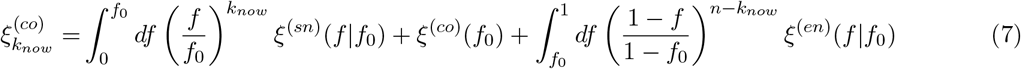

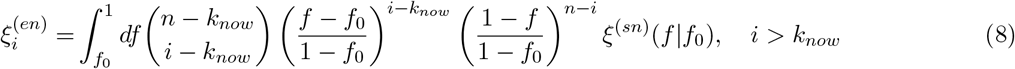

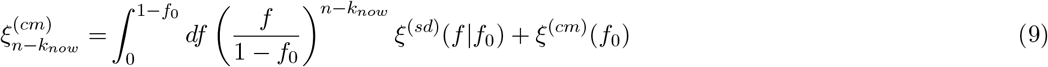

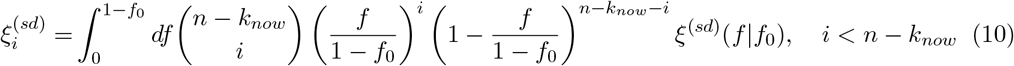

where 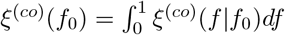 and 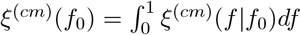.

Therefore the full spectrum for a rapid partial sweep is

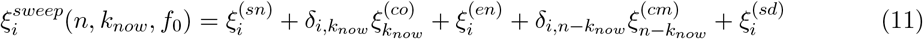

Finally, the contribution of the selected mutation should be added too (i.e., 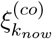 and 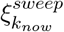 should be increased by 1).

### 2.5 Polygenic adaptation as a small sweep limit

Polygenic adaptation corresponds to many loci each undergoing small allele-frequency changes. In our framework, this is the limit where the derived focal allele frequency changes by a small amount

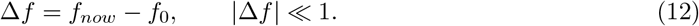

In this regime, the transformation of the SFS can be linearized, yielding an explicit first-order distortion around neutrality.

The distortion is basically undetectable locally, but it can leave a footprint across the genome. Assume in a genome of length *L*, there is a (small) probability *p*_*ps*_(*s*) that a mutation at any site would have a selection coefficient *s* under polygenic selection. Each of these sites is typically linked to *L*_*linked*_ sites. The distribution of mutational frequencies for these sites will be the neutral one, i.e. *θ ds p*_*ps*_(*s*)*/f*. The shift in frequency for a mutation of frequency *f* and selection coefficient *s* will be Δ*f* = *sf* (1 − *f*)*t*_*s*_. Hence, the distortion of the spectrum will be

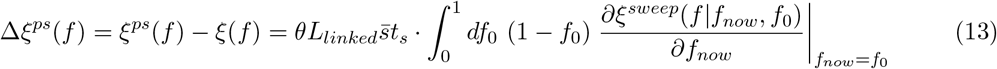

where 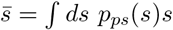 is the average fitness effect under polygenic selection.

## 3 Results

### 3.1 Local SFS distortion under polygenic selection

In the small-shift regime, the SFS under polygenic selection is

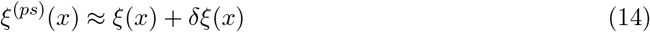

To connect this distortion to a biologically interpretable selection intensity, we consider a simple model where selected alleles are initially distributed neutrally and the expected frequency change scales as Δ*f* = *s f* (1 − *f*) *t*_*s*_, where *s* captures the strength of selection under the new environment and *t*_*s*_ the duration of selection. Under these assumptions, we provide an explicit expression for *δξ*(*x*) as a function of *x*, capturing how allele-frequency shifts at many loci collectively bias the distribution of linked variants:

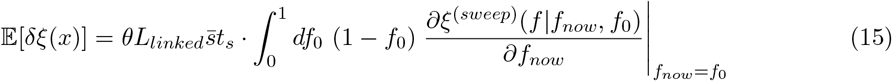

where the expected spectrum *ξ*^(*sweep*)^ is shown in the next section.

As expected, the magniture of this distortion depends on the typical linkage distance, the average selection strength for polygenic selection, and the time elapsed since the environmental change.

### 3.2 Spectrum under strong selection on an individual locus

For strong selection on an individual locus (rapid sweep from standing variation), we provide a simple analytical form for the components of the expected population SFS in the no-recombination limit:

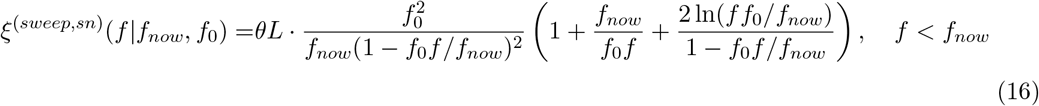

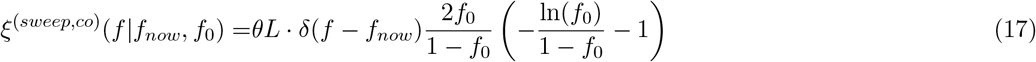

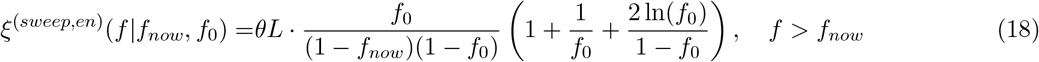

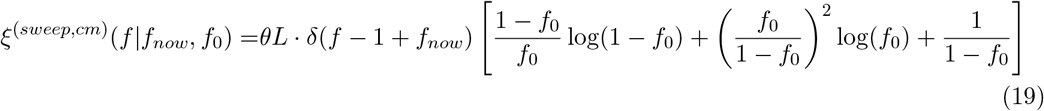

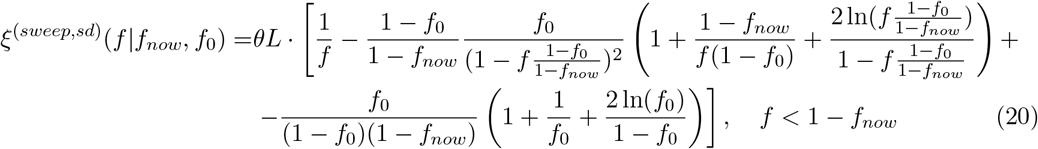

The spectrum is given by the sum of all these components:

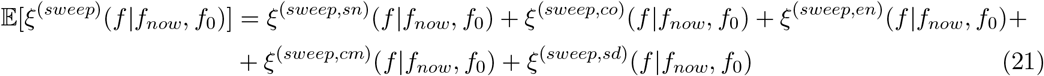

The discrete spectrum is discussed in the Methods and illustrated in Figures 1 in comparison with the classical neutral spectrum. Note that the presence of a linked neutral variant alone influences the expected spectrum, and therefore can be an appropriate comparison in some scenarios. We show the deviations caused purely by the strong selection in Figure 2.

**Figure 1.**
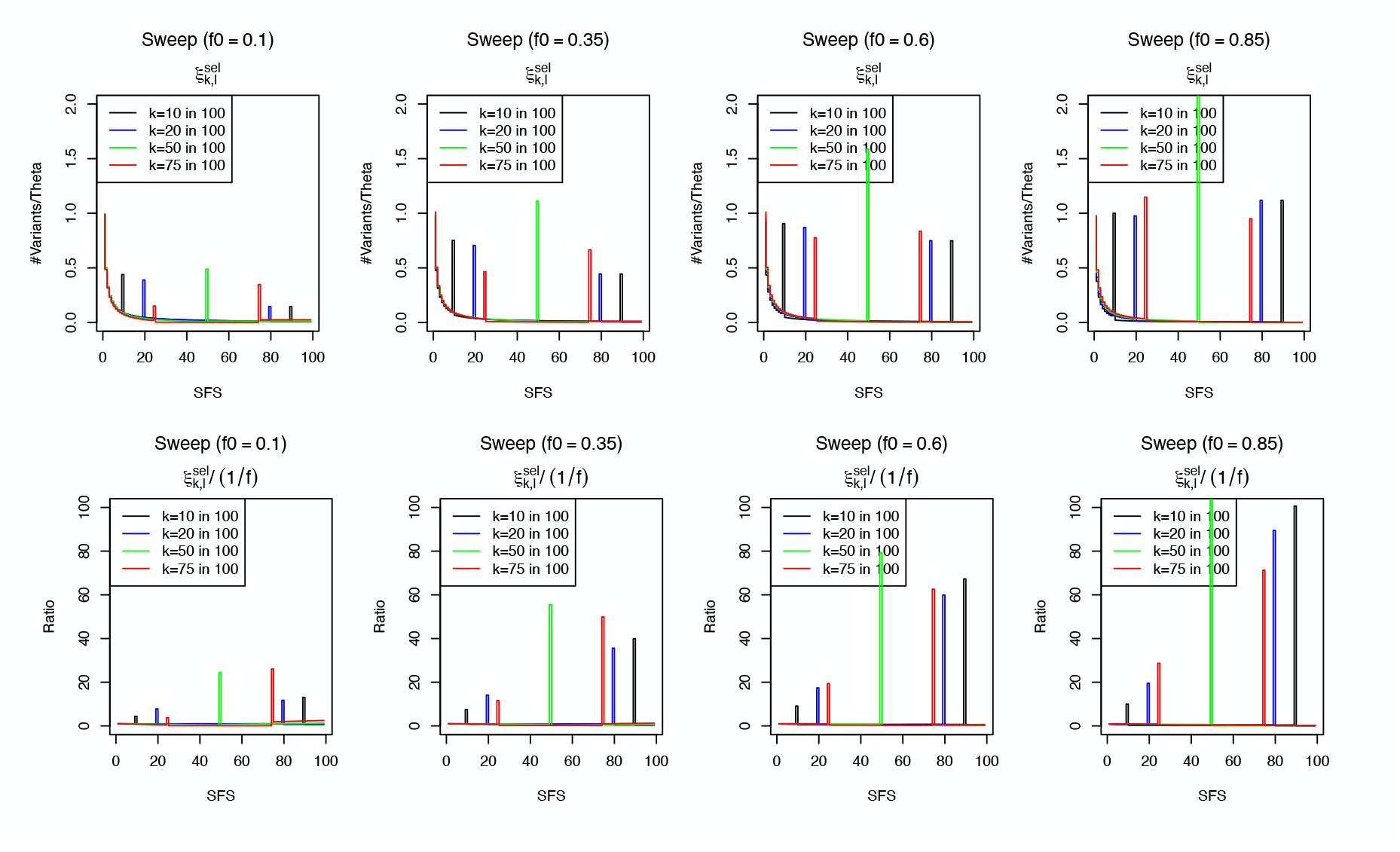
SFS after environmental change drives a rapid partial sweep from standing variation. Discrete spectrum 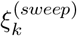 (first row) for a sample of size 100, and discrete spectrum compared to the neutral baseline *ξ*_*k*_ (second row), for different initial (*f*_0_) and final (*k*) frequencies

**Figure 2.**
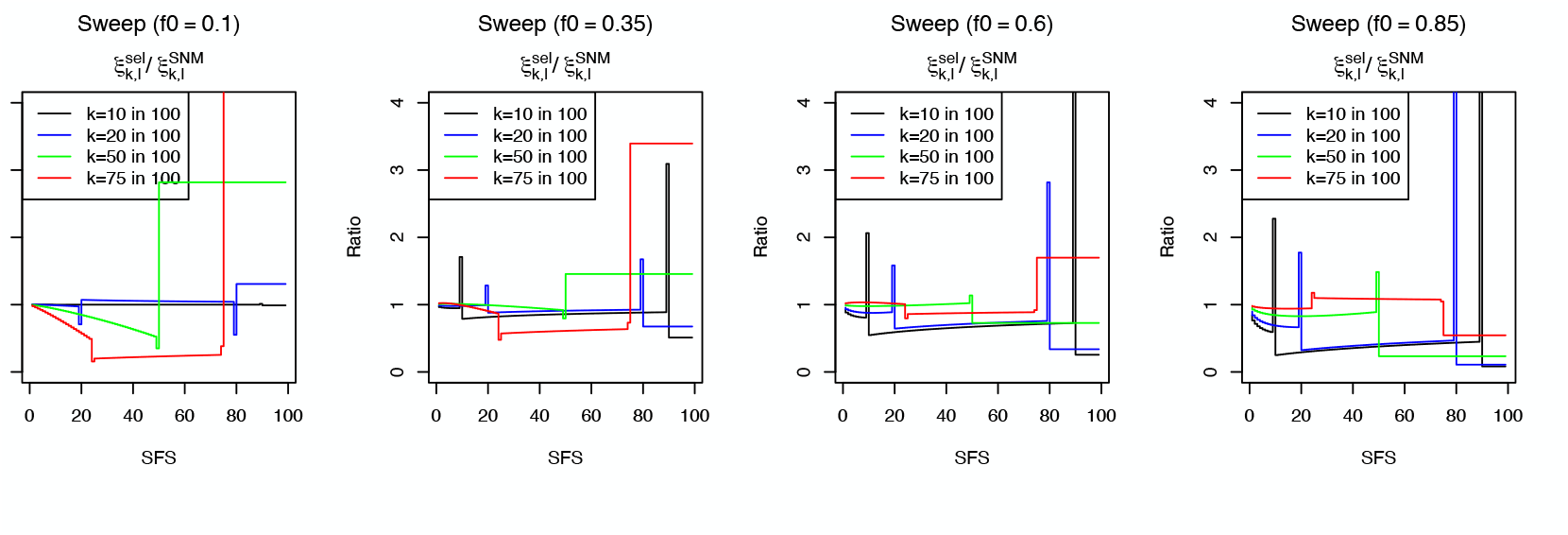
SFS after environmental change drives a rapid partial sweep from standing variation, compared with SFS linked to a neutral variant. Ratio of discrete spectra 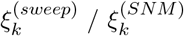 for different initial (*f*_0_) and final (*k*) frequencies.

### 3.3 Impact on genetic diversity statistics

From the transformed spectrum, we compute expectations for common diversity estimators, including Watterson’s estimator 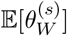 and nucleotide diversity 𝔼 [*π*^(*s*)^]. Because these statistics weight frequency classes differently, their relative changes provide a compact summary of the sweep- or shift-induced distortion. The results are shown in Figures 3 compared to neutral expectations, while the impact of rapid selection is illustrated in Figure 4. It is clear that the biggest impact of selection is a strong depletion of diversity for the cases closer to classical sweeps, i.e. for strong selection on rare alleles bringing them close to fixation.

**Figure 3.**
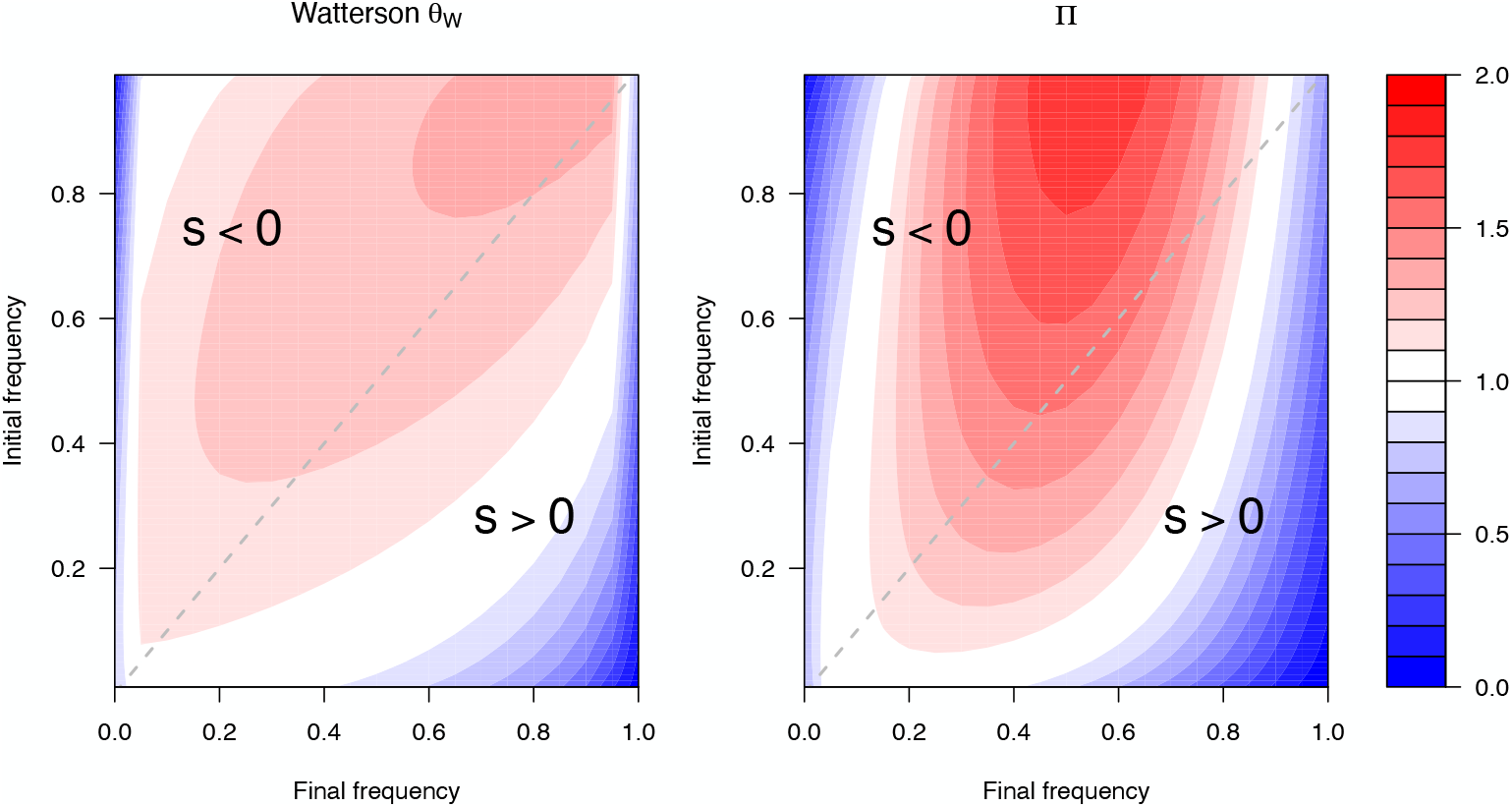
Predicted changes in diversity near loci involved in rapid adaptation. Normalized diversity summaries (e.g. *θ*_*W*_ */θ*_*W*,0_ and *π/π*_0_) as a function of initial and final derived allele frequencies.

**Figure 4.**
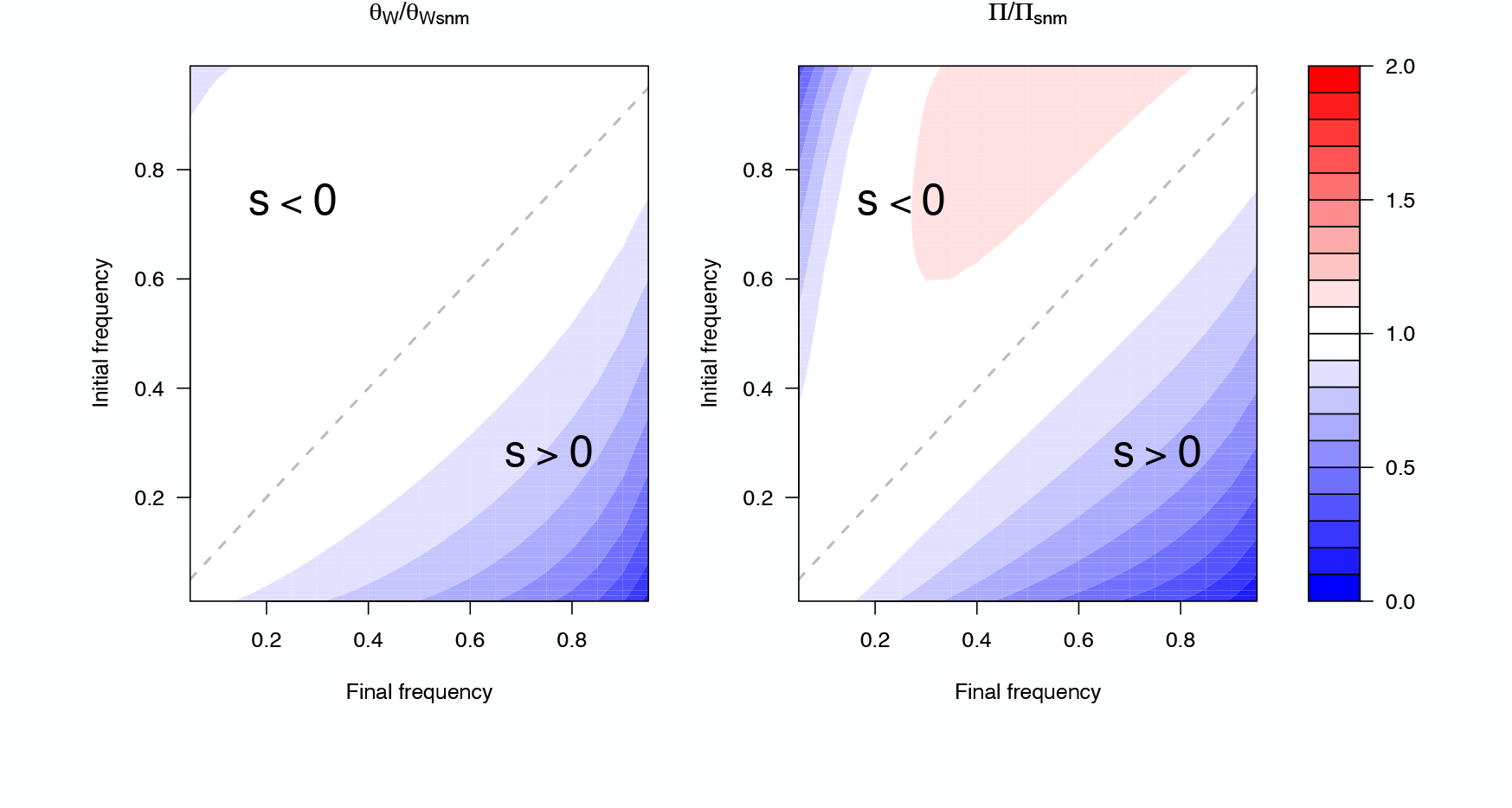
Predicted changes in diversity near loci involved in rapid adaptation vs near a neutral variant. Normalized diversity summaries (e.g. *θ*_*W*_ */θ*_*W,SNM*_ and *π/π*_*SNM*_) as a function of initial and final derived allele frequencies.

### 3.4 Impact on neutrality tests

We next evaluate the expected behavior of classical neutrality tests, including Tajima’s *D* and Fay and Wu’s *H*, under our rapid-adaptation model. These tests are usually centered (i.e. $*mathbbE*[*D*^(*s*)^] = 𝔼 [*H*^(*s*)^] = 0) and they are sensitive to different aspects of SFS shape; thus, their joint behavior can help distinguish rapid selection from demographic changes or other violations of the standard neutral model, especially when signals are localized around candidate loci.

The expected shifts in the average Tajima’s *D* and Fay and Wu’s *H* are shown in Figures 5 and 6 respectively. The impact of strong selection alone is shown in Figures 7 and 8, painting a complex picture. Both these tests show a strong negative bias for rare selected variants which undergo a sizable sweep. Tajima’s *D* values are however clearly inflated when a rare ancestral variant is undergoing a partial sweep that brings it to intermediate frequencies.

**Figure 5.**
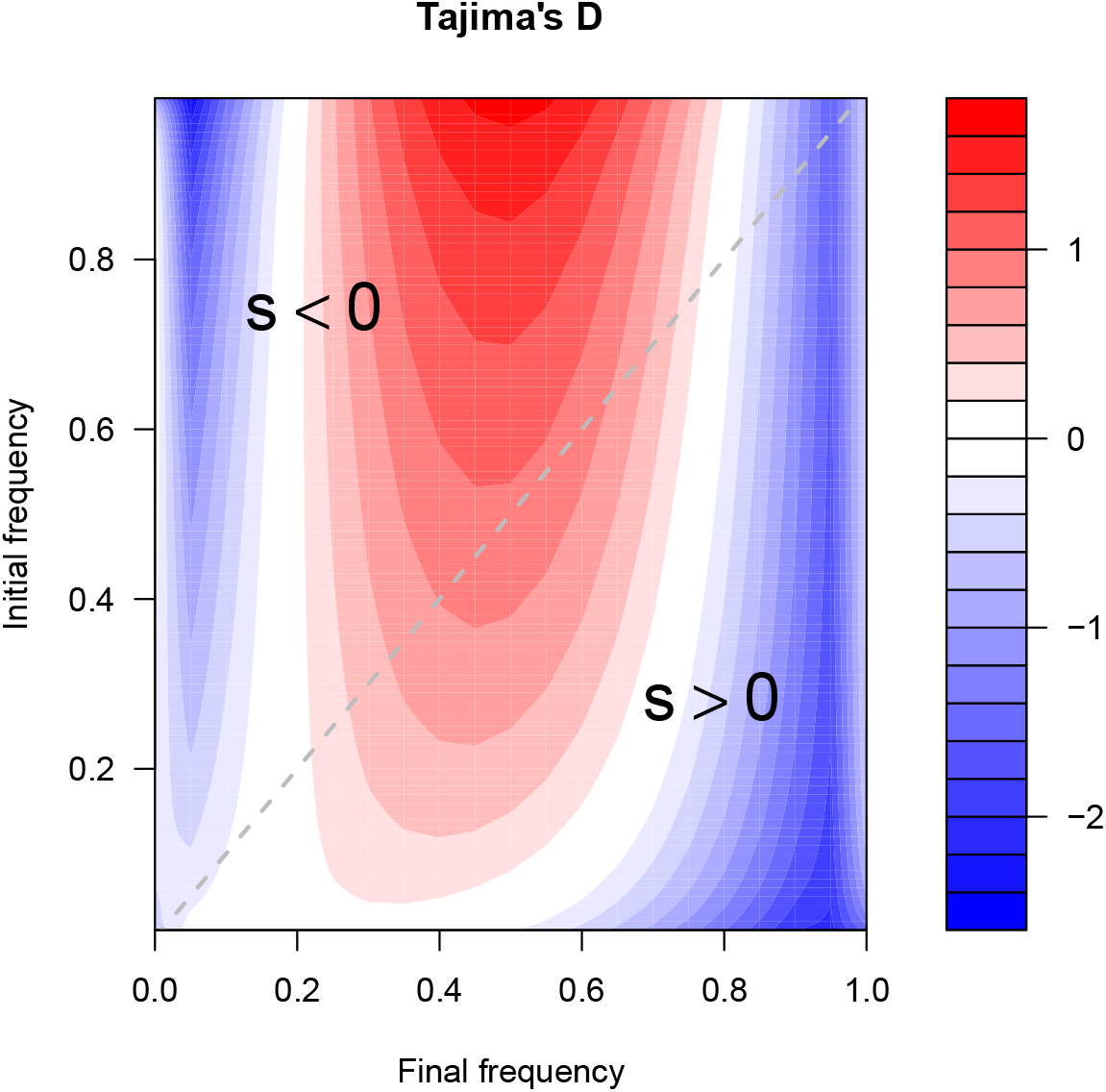
Predicted shifts in neutrality statistics near rapidly adapting loci. Expected Tajima’s *D* of initial and final derived allele frequencies, compared to the neutral baseline.

**Figure 6.**
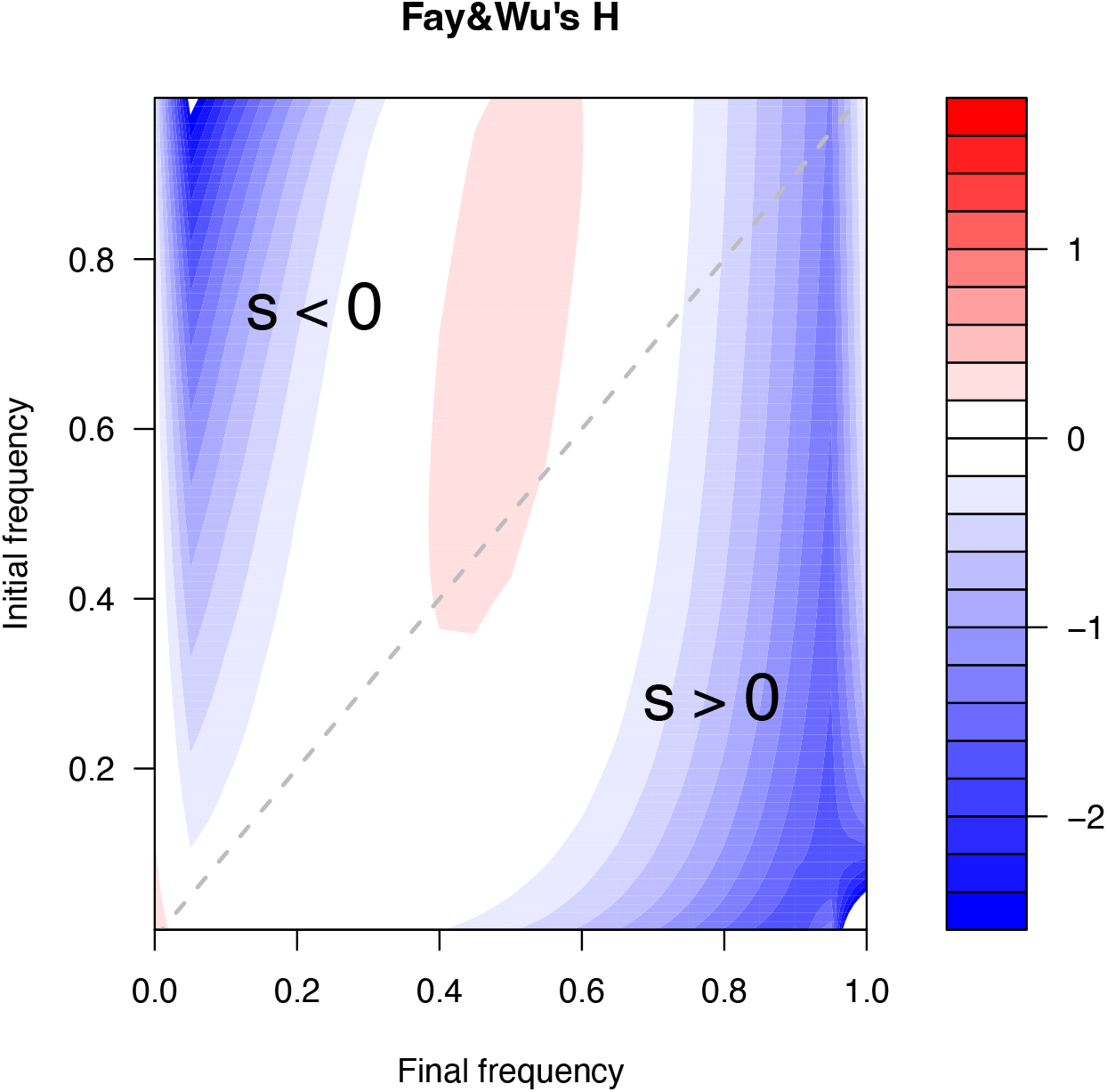
Predicted shifts in neutrality statistics near rapidly adapting loci. Expected Fay and Wu’s *H* as a function of initial and final derived allele frequencies, compared to the neutral baseline.

**Figure 7.**
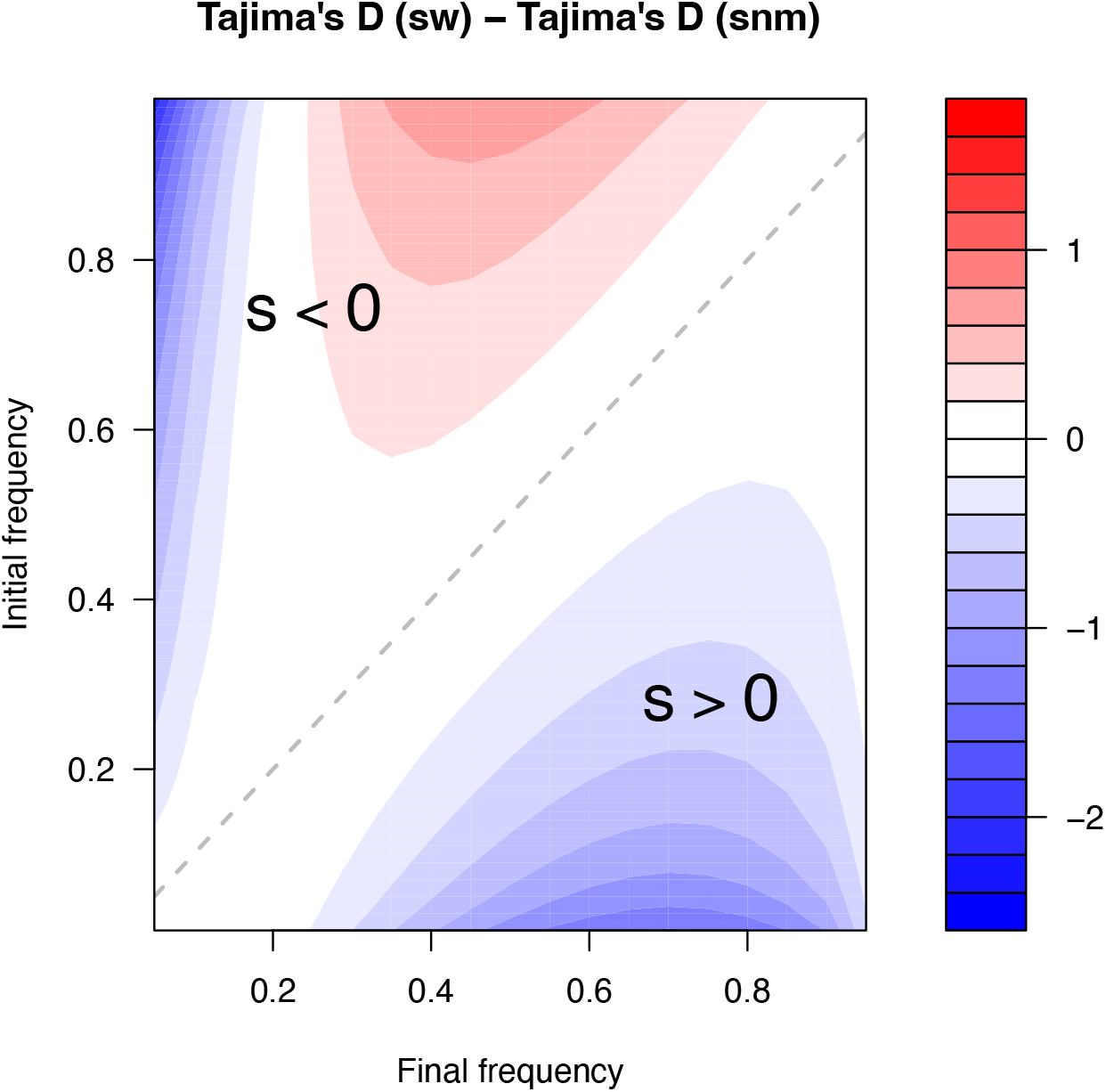
Predicted shifts in neutrality statistics near rapidly adapting loci vs near a neutral variant. Expected Tajima’s *D* as a function of initial and final derived allele frequencies, compared to a conditional neutral baseline.

**Figure 8.**
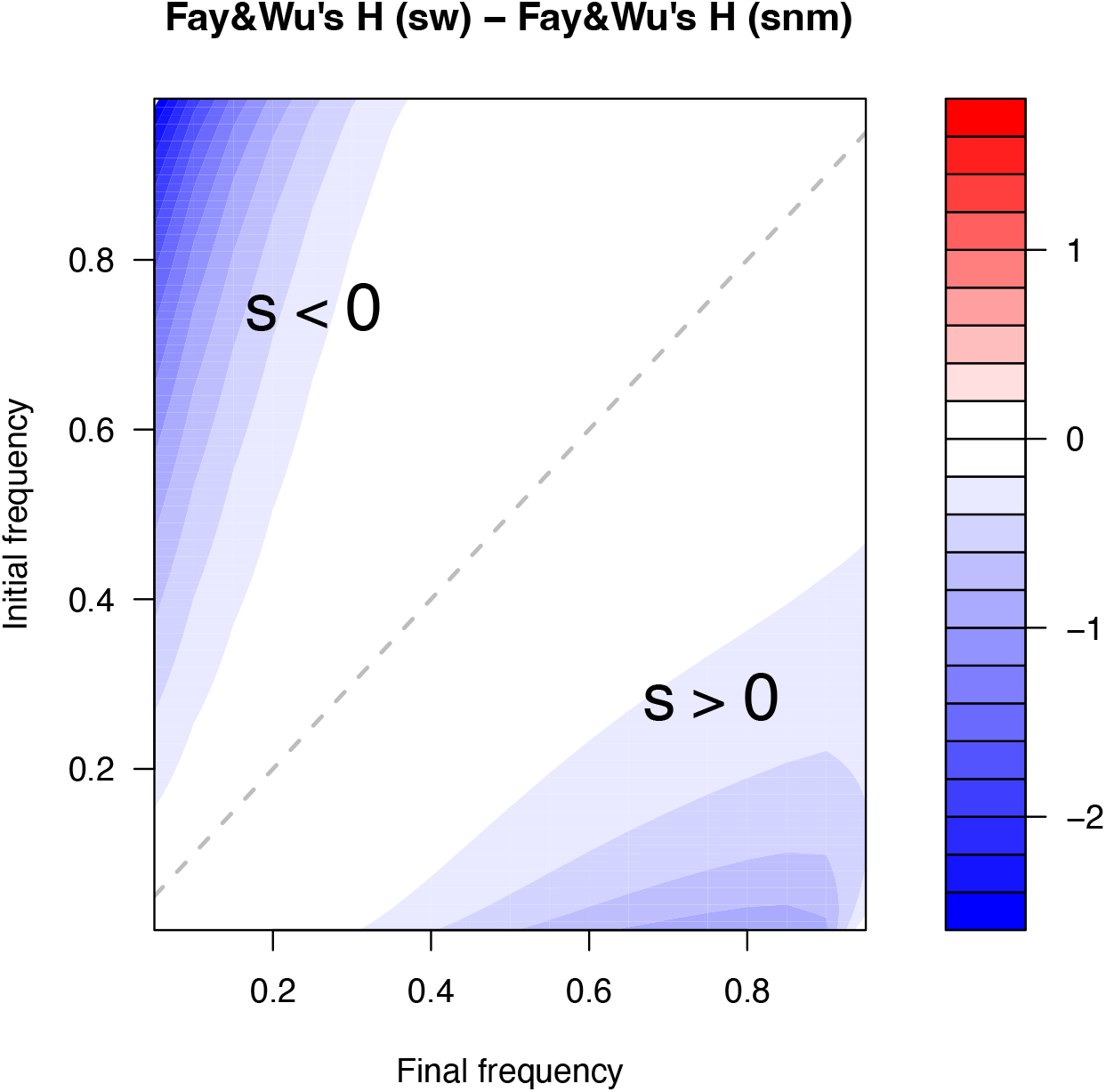
Predicted shifts in neutrality statistics near rapidly adapting loci vs near a neutral variant. Expected Fay and Wu’s *H* as a function of initial and final derived allele frequencies, compared to a conditional neutral baseline.

## 4 Discussion

We presented explicit analytical results for the genetic footprints expected when populations adapt rapidly to sudden, extreme, and unprecedented environmental change. Our framework captures two limiting modes: (i) single-locus adaptation (rapid sweep from standing variation), which produces strong localized distortion of linked variation near the selected site; and (ii) polygenic adaptation (many small allele-frequency shifts), which produces weaker but systematic distortions that can accumulate across loci and can be summarized by first-order deviations from neutrality.

Which regime dominates in practice depends on trait architecture, the distribution of effect sizes, the amount of standing variation, and the selection pressures imposed by the new environment Wilson *et al*. (2017). In small populations, rapid adaptation often requires either alleles already segregating at appreciable frequencies or variants of sufficiently large effect that selection can overcome drift on short timescales. Discriminating between all these signatures is notoriously difficult Schrider *et al*. (2015).

A primary limitation of our analysis is the focus on the region closest to the selected variant, where recombination during the short adaptive episode can be ignored. At larger genetic distances, recombination will partially uncouple linked variants from the selected background, altering the initial pre-selection linked spectrum and changing how the post-selection rescaling acts Berg and coop (2015). A simulation-based treatment of recombination of these rapid-adaptation signatures has been recently presented in Osmond and coop (2020).

We assumed that linked variants were neutral before the environmental shift. Many natural populations harbor nearly neutral variants whose dynamics before the shift are shaped by weak selection and drift. Because our key results is driven by rapid changes in proportions of genetic backgrounds during the adaptive episode, we expect qualitatively similar distortions for almost-neutral variation in the immediate vicinity of the selected site.

Finally, standing genetic variation is not the only substrate for rapid response. Adaptation may also proceed via de novo mutations of large effect (including structural variants), migration and genetic rescue through crossing between populations, and introgression from other species or divergent lineages. These processes can generate footprints that overlap with (or differ from) the patterns described here, particularly when introgressed haplotypes carry blocks of linked variation that mimic sweep-like signals. Integrating these routes into a unified framework represents an important direction for future work.

